# Loss of ZnT8 function protects against diabetes by enhanced insulin secretion

**DOI:** 10.1101/436030

**Authors:** Om Prakash Dwivedi, Mikko Lehtovirta, Benoit Hastoy, Vikash Chandra, Sandra Kleiner, Deepak Jain, Ann-Marie Richard, Nicola L. Beer, Nicole A. J. Krentz, Rashmi B. Prasad, Ola Hansson, Emma Ahlqvist, Ulrika Krus, Isabella Artner, Daniel Gomez, Aris Baras, Fernando Abaitua, Benoite Champon, Anthony J Payne, Daniela Moralli, Soren K. Thomsen, Philipp Kramer, Ioannis Spiliotis, Reshma Ramracheya, Pauline Chabosseau, Andria Theodoulou, Rebecca Cheung, Martijn van de Bunt, Jason Flannick, Maddalena Trombetta, Enzo Bonora, Claes B. Wolheim, Leena Sarelin, Riccardo C. Bonadonna, Patrik Rorsman, Guy A Rutter, Benjamin Davies, Julia Brosnan, Mark I. McCarthy, Timo Otonkoski, Jens O. Lagerstedt, Jesper Gromada, Anna L. Gloyn, Tiinamaija Tuomi, Leif Groop

## Abstract

A rare loss-of-function variant p.Arg138* in *SLC30A8* encoding the zinc transporter 8 (ZnT8) enriched in Western Finland protects against type 2 diabetes (T2D). We recruited relatives of the identified carriers and showed that protection was associated with better insulin secretion due to enhanced glucose responsiveness and proinsulin conversion, especially compared with individuals matched for the genotype of a common T2D risk variant in *SLC30A8*, p.Arg325. In genome-edited human IPS-derived β-like cells, we establish that the p.Arg138* variant results in reduced *SLC30A8* expression due to haploinsufficiency. In human β-cells loss of *SLC30A8* leads to increased glucose responsiveness and reduced K_ATP_ channel function, which was also seen in isolated islets from carriers of the T2D-protective allele p.Trp325. These data position ZnT8 as an appealing target for treatment aiming at maintaining insulin secretion capacity in T2D.

## Introduction

Zinc transporters (ZnTs) regulate the passage of zinc across biological membranes out of the cytosol, while Zrt/Irt-like proteins transport zinc into the cytosol^1^. ZnT8, encoded by *SLC30A8*, is highly expressed in membranes of insulin granules in pancreatic β-cells, where it transports zinc ions for crystallization and storage of insulin^2^. We have described a loss-of-Function (LoF) variant p.Arg138* (rs200185429, c.412C>T) in the *SLC30A8* gene, which conferred 53% protection against T2D^3^. This variant was extremely rare (0.02%) in most European countries but more common (>0.2%) in Western Finland^3^. We also reported a protective frameshift variant p.Lys34Serfs*50 conferring 83% protection against T2D in Iceland. A recent (>44K) exome sequencing study reported >30 alleles in *SLC30A8* reducing the risk of T2D, confirming it as a robust target for T2D protection^4^. Further, the *SLC30A8* gene also harbors a common variant (rs13266634, c.973T>A) p.Trp325Arg in the C-terminal domain^5^. While the major p.Arg325 allele (>70% of the population) confers increased risk for T2D, the minor p.Trp325 allele is protective^6^.

The mechanisms by which reduced activity of ZnT8 protect against T2D are largely unknown. Several attempts have been made to study loss of *Slc30a8* function in rodent models, but the results have been inconclusive: knock-out of *Slc30a8* led to either glucose intolerance or had no effect in mice^7, 8, 9^, while over-expression improved glucose tolerance without effect on insulin secretion^10^. In a mouse model harbouring the equivalent of the human p.Arg138* variant we were unable to detect any ZnT8 protein and observed no effect on glucose^11^. These rodent *in vitro* and *in vivo* experiments present a complex picture which might not recapitulate the T2D protective effects by *SLC30A8* LoF mutations in humans. We therefore performed detailed metabolic studies in human carriers of the LoF variant (p.Arg138*) recruited on the basis of their genotype, performed comprehensive functional studies in human β-cell models and compared with the mouse model carrying the human p.Arg138*-*SLC30A8* mutation.

## Results

### Recruitment by genotype

Given the enrichment of the p.Arg138\**-SLC30A8* variant in Western Finland, we genotyped >14,000 individuals from the Botnia Study^12^ for the *SLC30A8* p.Arg138* mutation and the common p.Trp325Arg variant (Fig. 1). None of the p.Arg138* mutation carriers was homozygous for the protective common variant, p.Trp325 and p.Arg138* segregated with p.Arg325 in the families (Supplementary Fig. 1). Thus, we present the data in three different ways: 1) p.Arg138* *vs.* all p.Arg138Arg, 2) p.Arg138* *vs.* p.Arg138Arg having at least one p.Arg325 allele (p.Trp325Arg or p.Arg325Arg), and 3) p.Arg325 (p.Trp325Arg or p.Arg325Arg) *vs.* p.Trp325Trp on a background of p.Arg138Arg. We included 79 p.Arg138* carriers and 103 non-carriers. Of them, 54 p.Arg138* and their 47 relatives with p.Arg138Arg participated in a test meal (Fig. 1 and Supplementary Table 1). In addition, 35 p.Arg138* and 8141 p.Arg138Arg had previously undergone an oral glucose tolerance test (OGTT, Fig. 1 and Supplementary Table 2).

**Fig. 1:**
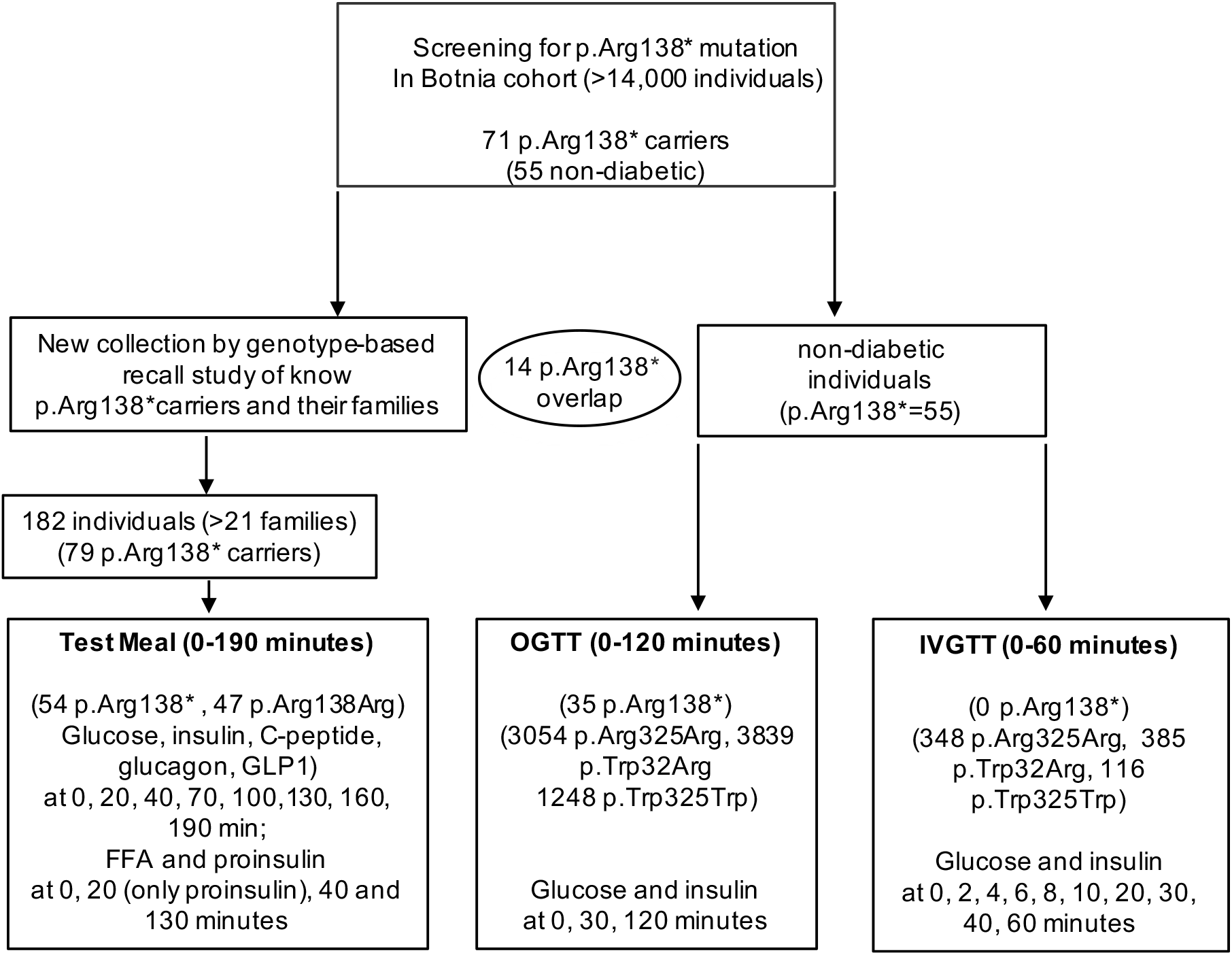
A flow-chart of study design for human *in vivo* studies

Replicating our previous findings^3^, carriers of p.Arg138* had reduced risk of T2D (OR=0.40, P=0.003) in analysis of total 4564 T2D subjects (13 p.Arg138* carriers) and 8183 non-diabetic (55 p.Arg138* carriers) individuals. Additionally, non-diabetic p.Arg138* carriers have lower fasting glucose concentrations (P=0.033) than p.Arg138Arg. There were no significant differences in plasma zinc concentrations measured during test meal or OGTT (data not shown).

#### Comparison of p.Arg138* vs. p.Arg138Arg

The p.Arg138* carriers have lower blood glucose levels during test meal specifically during the first 40 minutes (P=0.02) and better corrected insulin response (CIR) (at 20 min, p=0.046) than non-carriers (Fig. 2a and Supplementary Tables 3). Similarly, the carriers had better insulin response to OGTT (Fig 3b-c, left panel), especially the early incremental insulin response (p=0.008) and insulin/glucose ratio (at 30 min, p=0.002, Supplementary Tables 4). Of note, the p.Arg138* carriers had significantly lower proinsulin/C-peptide (20 min: P=0.041; 40 min: P=0.043) and proinsulin/insulin (20 min: P=0.006) ratios during test meal suggesting effects on proinsulin conversion (Fig. 2d-e). No differences were seen in glucagon, GLP-1 or free fatty acids concentrations during test meal (Supplementary Fig. 2c-e). Neither model-based insulin clearance index nor the ratio of insulin and C-peptide areas under the curve during test meal differed between p.Arg138* and p.Arg138Arg, making changes in insulin clearance^13^ unlikely (Supplementary Fig. 2f-g).

**Fig. 2:**
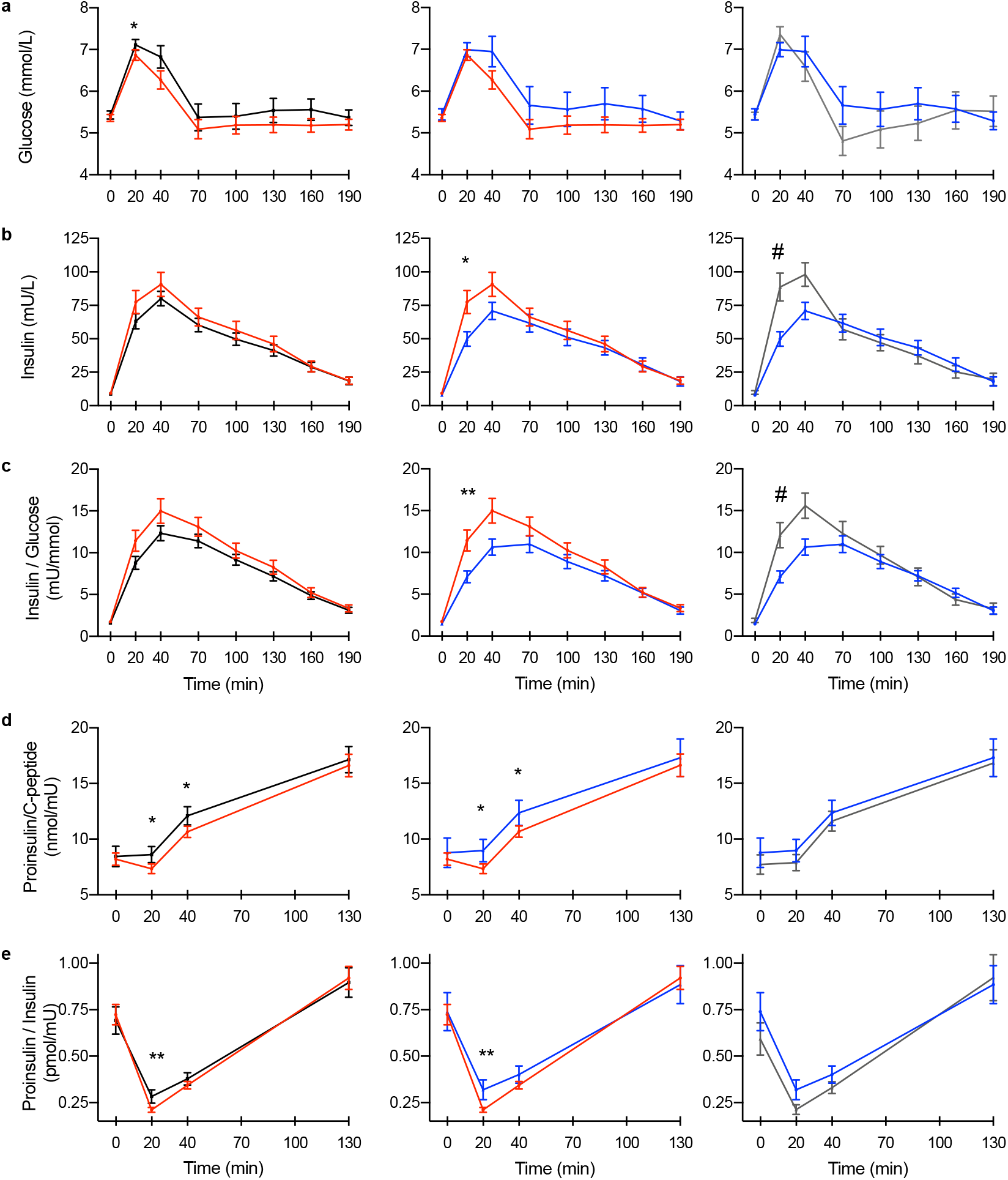
SLC30A8-p.Arg138* enhances insulin secretion and proinsulin processing during test meal. Association of *SLC30A8* p.Arg138* and p.Trp325Arg variants with **a,** plasma glucose **b,** serum insulin **c,** insulin/glucose ratio **d,** proinsulin/C-peptide ratio and **e,** proinsulin/insulin ratio during test meal. *Left panel*: Carriers (red, N=54) vs. non-carriers (black, N=47) of p.Arg138*. *Middle panel*: Carriers of p.Arg138* (red, N=54) vs Arg138Arg having the common risk variant p.Arg325 (blue, N=31). *Right panel*: Carriers of p.Trp325Trp (grey, N=16) vs. p.Arg325 (blue, N=31). Data are Mean ± SEM. P-values were calculated by family-based association (*) or linear regression (#) (adjusted for age, sex, BMI and p.Trp325Arg variant status for the middle pane, Methods): */#, p < 0.05, **/##, p < 0.01.

**Fig. 3:**
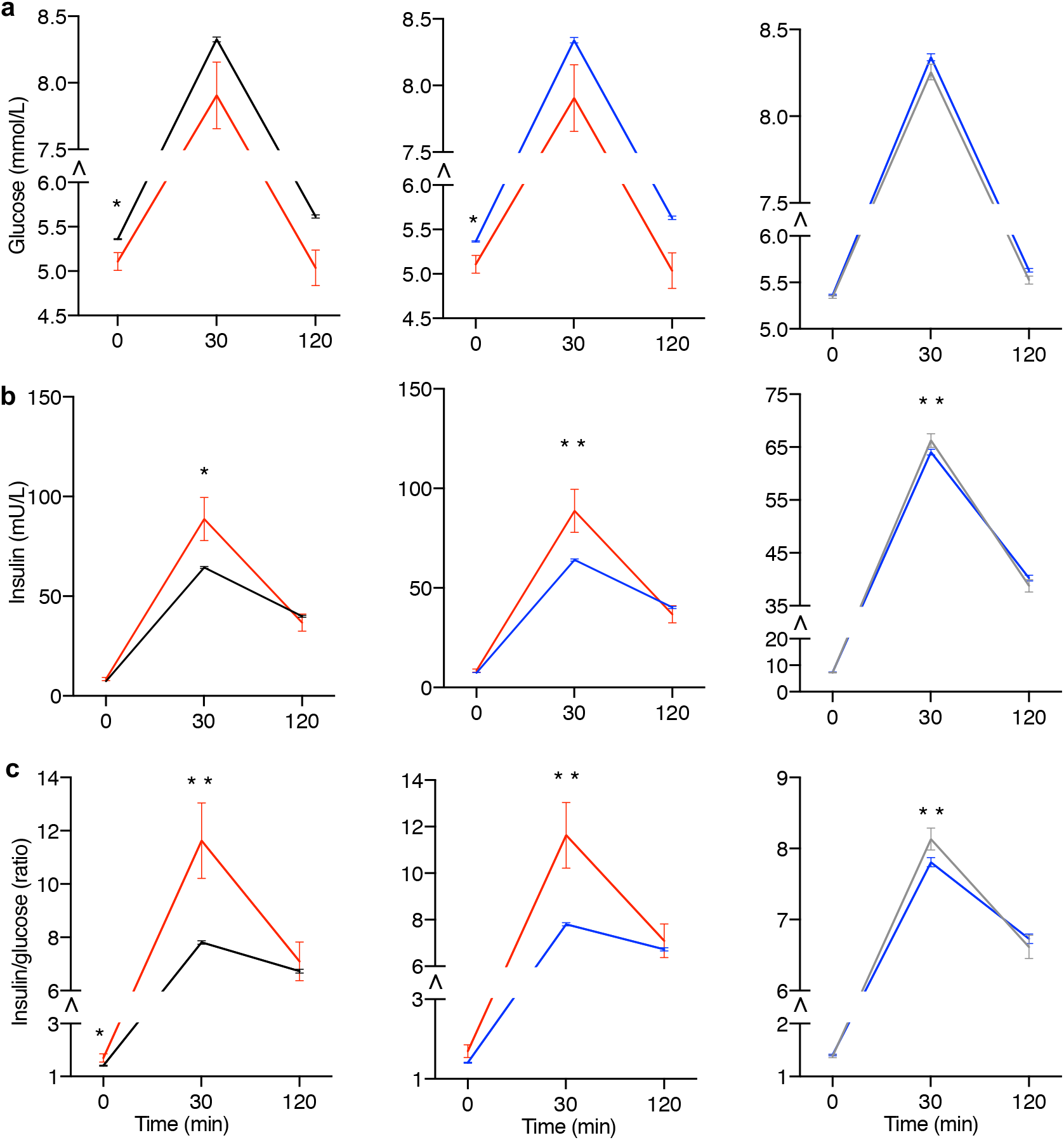
*SLC30A8* p.Arg138* and p.Trp325 enhance insulin secretion during OGTT. Association of *SLC30A8* p.Arg138* and p.Trp325Arg with **a**, plasma glucose **b**, serum insulin **c**, insulin/glucose ratio during an oral glucose tolerance test (OGTT). *Left panel*: Carriers (red, N=35) vs. non-carriers (black, N=7954-8141) of p.Arg138*. *Middle panel*: Carriers of p.Arg138* (red, N=35) vs. p.Arg138Arg having the common risk variant p.Arg325 (blue, N=6728-6893). *Right panel*: Carriers of p.Trp325Trp (grey N=1226-1248) vs. p.Arg325 (blue, N=6728-6893). Data are shown as Mean ±SEM. P-values (mixed model, Methods) using additive effect: * < 0.05, ** < 0.01. Y-axis: note truncation (⋀) and different scale in the right panel.

#### Comparison of p.Arg138* vs. p.Arg138Arg–p.Arg325

The above differences were magnified when we restricted the p.Arg138Arg group to carriers of the common risk variant p.Arg325 (middle panel of Fig. 2). The early phase (0-40 min) insulin (p=0.026), insulin/glucose ratio (p=0.004) and CIR (p=0.004; 20 min, Supplementary Table 3) were all greater in p.Arg138* carriers compared with those having p.Arg138Arg on a background of p.Arg325. Both the proinsulin/C-peptide (20 min: P=0.027, 40 min: P=0.044) and proinsulin/insulin ratios (20 min: P=0.003) were reduced in p.Arg138* carriers (middle panel of Fig. 2d-e).

#### Comparison of p.Trp325Trp vs. p.Arg325

The effect of p.Trp325Trp genotype on glucose and insulin response mimicked the effects of p.Arg138* with pronounced early (20 min) insulin (p=0.035) and C-peptide (p=0.025) responses during test meal (right panel of Fig. 2b-c and Supplementary Fig. 2a), as well as increased insulin secretion (30 min insulin, 30 min insulin/glucose, incremental insulin, P≤0.003) and lower fasting and 120 minute proinsulin (p=0.006 and p=0.039, respectively) concentration during OGTT in p.Trp325 carriers (Supplementary Table 4, right panel of Fig. 3b-c). Moreover, p.Trp325Trp carriers undergoing intravenous glucose tolerance tests (IVGTT) showed a pronounced (p=0.003) early incremental insulin secretion response (Supplementary Fig. 3a-b and Supplementary Table 4). In patients with newly diagnosed T2D, the p.Trp325Trp carriers showed a trend (P=0.12) to enhanced β-cell sensitivity to glucose during the OGTT (Supplementary Fig. 3c).

Taken together, all the human *in vivo* results show that T2D protection by the LoF variant p.Arg138* is due to enhanced glucose-stimulated insulin secretion combined with enhanced proinsulin conversion. The common T2D protective allele p.Trp325 shows a similar – albeit weaker - metabolic phenotype suggesting it might also reduce ZnT8 function.

### *SLC30A8* p.Arg138* variant in human iPSCs

The majority of nonsense *SLC30A8* alleles (including p.Arg138*) protecting against T2D are located in the first four exons of the eight-exon canonical islet *SLC30A8* transcript ENST00000456015 and are predicted to undergo nonsense mediated decay (NMD), a cell surveillance pathway which reduces errors in gene expression by eliminating mRNA transcripts that contain premature stop codons. To confirm that the p.Arg138* allele indeed leads to haploinsufficiency through NMD, we used CRISPR-Cas9 to introduce the p.Arg138* variant into the *SLC30A8* locus of the SB Ad3.1 human iPSC cell line (Supplementary Fig. 4a, Methods). Two hiPSC lines for the p.Arg138*-SLC30A8 variant (Clone B1 and A3) were generated and compared to an unedited p.Arg138Arg-SLC30A8 CRISPR hiPSC line. Both B1 and A3 clones were heterozygous with mono-allelic sequencing confirming the p.Arg138* variant in only one allele (Supplementary Fig. 4b). All hiPSC lines passed quality control checks including karyotyping and pluripotency (Supplementary Fig. 4c).

**Fig. 4:**
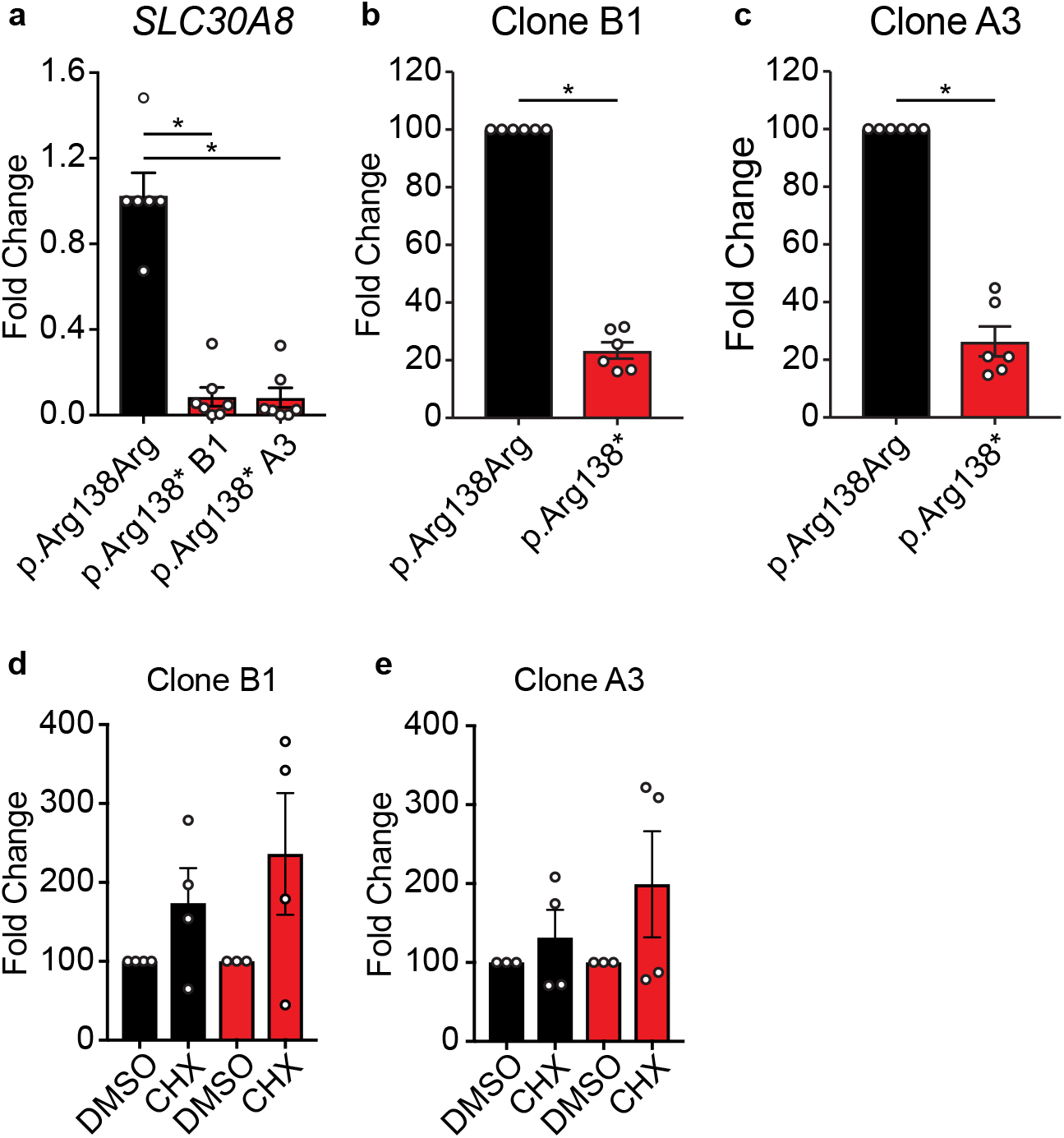
Beta like cells derived from SLC30A8-p.Arg138* iPSCs display haploinsufficiency of *SLC30A8*. **a**, *SLC30A8* expression in cells heterozygous for *SLC30A8-*p.Arg138*. Data normalized to *TBP gene* are expressed as fold change relative to p.Arg138Arg control (n=6-7 wells from two differentiations). Allele-specific expression (ASE) of p.Arg138Arg (black bar) and p.Arg138* (red bar) in **b**, clone B1 or **c**, clone A3 derived cells. Allele-specific expression of p.Arg138Arg (black bar) and p.Arg138* (red bar) in **d**, clone B1 and **e**, clone A3 derived cells treated with DMSO (Dimethyl sulfoxide) or cycloheximide (CHX) for four hours. ASE data (Mean ±SEM) were determined by Digital Droplet PCR and presented as fold change relative to p.Arg138Arg transcript **(b, c,** n=6 wells from two differentiations) or to DMSO control (**d-e,** n=3-4 wells from two differentiation). * P<0.05 (Kruskal-Wallis test for multiple comparisons or unequal variance t-test).

Accordingly, we subjected our *SLC30A8-*edited iPSCs to a previously published *in vitro* endocrine pancreas differentiation protocol^14^ (Supplementary Fig. 4d-k, Methods). At the end of the seven stage protocol, *SLC30A8* expression was significantly reduced in cells heterozygous for the p.Arg138* allele (clone B1 0.09±0.04; clone A3 0.08±0.05) compared to unedited control cells (1.03±0.11) (Fig. 4a). Of note, p.Arg138* allele specific *SLC30A8* expression was reduced compared to the WT allele^15^ (clone B1: 22.9±2.1%; clone A3: 26.0±3.9%) (Fig. 4b-c). Inhibition of NMD by cyclohexamide increased expression of the p.Arg138* transcript more than the p.Arg138Arg transcript compared to DMSO control (clone B1:209±52% and clone A3: 199±67% *vs*. clone B1: 161±30% and clone A3: 132±35%, respectively, Fig. 4d-e). Taken together, these data show that the protective p.Arg138*-*SLC30A8* allele undergoes NMD, resulting in haploinsufficiency for *SLC30A8*.

### Impact of *SLC30A8* loss in a human β-cell line

Since human *in vivo* studies provided strong evidence for a role of the p.Arg138* on insulin secretion and proinsulin processing, we studied the impact of *SLC30A*8 loss using siRNA mediated knock down (KD) on both phenotypes in a well characterized human β-cell model EndoC-βH1^16^ (Methods). By siRNA, we achieved 55-65% decrease in *SLC30A8* mRNA (p=0.008) and protein (p=0.016, Fig. 5a-c).

KD of *SLC30A8* had no significant effect on glucose-or tolbutamide-stimulated insulin secretion or on insulin content (Fig. 5d-e) but basal insulin secretion was higher in si*SLC30A8* transfected cells compared to scrambled siRNA cells (p=0.012, Fig. 5d), and the inhibitory effect of diazoxide, a K_ATP_ channel opener, on glucose-stimulated insulin secretion was reduced (p=2×10^-3^, Fig. 5d). We measured the resting membrane conductance (G_m_), which principally reflects K_ATP_ channel activity. In control cells, G_m_ was in agreement with that previously reported^17^. *SLC30A8* KD reduced G_m_ by 65% (p=0.002, Fig. 5f) without effect on cell size (Fig. 5g), an effect that correlated with reduced expression of the two genes encoding the K_ATP_ channel subunits SUR1 (*ABCC8)* and Kir6.2 (*KCNJ11)* (Fig. 5h). However, insulin secretion elicited by increasing extracellular K^+^ ([K^+^]_o_) to 50 mM (to depolarise the cells and open voltage-gated Ca^2+^ channels) and 16.7 mM glucose was significantly higher after *SLC30A8* KD (p=0.008, Fig. 5i). The proinsulin-insulin ratios (both total and secreted hormones) were decreased in si*SLC30A8* cells (p<0.001, Fig. 5j-k). Although mRNA of the proinsulin processing genes *PC1/3* and *CPE* was decreased, we could not detect a similar reduction at the protein level (Fig. 5l-n).

**Fig. 5:**
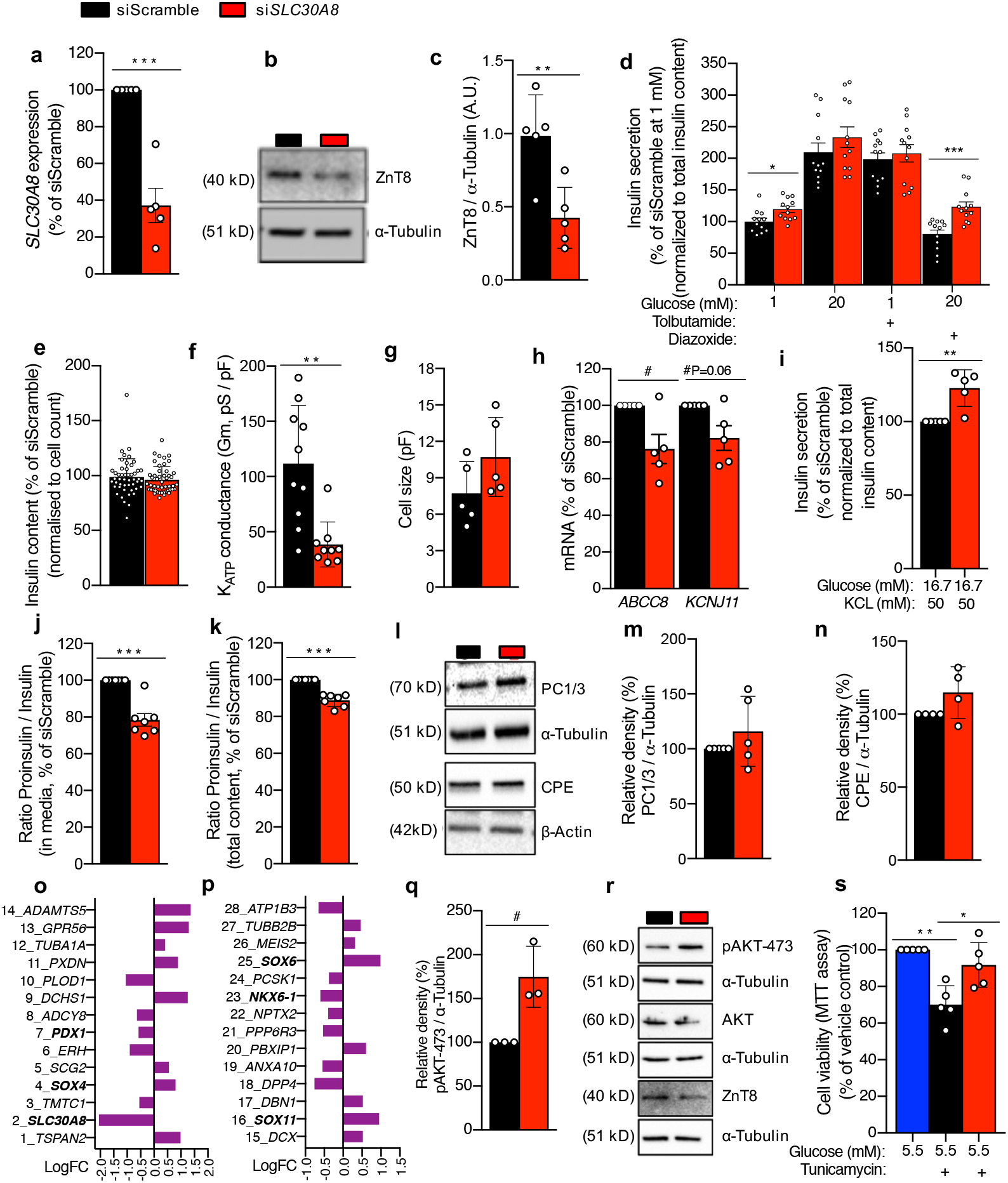
*SLC30A8* knock down leads to enhanced insulin secretion, proinsulin processing and cell viability in the human pancreatic EndoC-βh1 cells. **a-c,** Characterization of *SLC30A8* knock down (KD) at the (**a**) mRNA and protein level (**b**-immunoblot, **c**-densitometry). **d-i**, Effect of KD on (**d**) insulin secretion stimulated by glucose and K_ATP_ channel regulators (as labelled), (**e**) insulin content, (**f**) K_ATP_ channel conductance (Gm), (**g**) cell size, (**h**) expression of K_ATP_ channel subunits, (**i**) insulin secretion stimulated by KCL and high glucose. **j-n**, Effect of KD on proinsulin processing estimated by (**j-k)** proinsulin/insulin ratio and proinsulin processing enzymes PC1/3 and CPE **(l**, immunoblot, **m-n**, densitometry). **o-p**, Effect of KD (n=3 *vs.* 3) on whole transcriptome (mRNAs) by next generation sequencing and depicting 28 top candidate genes ranked by increasing p values (1% FDR corrected, P≤0.0002). **q-s**, Effect of KD on basal (5.5 mM glucose) AKT phosphorylation (**q**, densitometry, **r**, immunoblot; phospho-AKT-Ser473, total AKT) and cell viability under ER stress (**s**, MTT assay, 10 μg/ml tunicamycin, DMSO as vehicle control). Data are shown as Mean ±SEM (N=3-10). P-values (*Mann-Whitney test/#Unpaired t test): */# p ≤0.05, ** p < 0.01, *** p < 0.001.

RNA sequencing of *SLC30A8* KD cells (n=3 *vs.* 3) replicated the reduction of *KCNJ11* and *ABCC8* gene expression (p=4.3 ×10^-3^ and p=2.9×10^-5^, respectively). In addition, expression of genes involved in regulation of β-cell excitability was down-regulated, including *KCNMA1* encoding a Ca^2+^-activated K^+^ channel18 and *TMTC1* (p=6.8×10^-5^ and 2.9×10^-16^, respectively) encoding an ER adapter protein influencing intracellular calcium levels. Also, expression of genes associated with β-cell maturation and secretion was influenced by *SLC30A8* KD with decreased expression of *NKX6.1* and *PDX1* and increased expression of *SOX4*, *SOX6* and *SOX11* (Fig. 5o-p).

In addition, we also observed increased AKT phosphorylation (pAKT-473) and improved cell survival under ER stress (p<0.017, Fig. 5q-s), mechanisms which also could contribute to the overall protection by preserving β-cell mass^19^. Taken together, these data generated by disrupting *SLC30A8* in a human β-cell pointed at multiple mechanisms including changes in proinsulin conversion, K_ATP_ channel activity and cell viability.

### Metabolic phenotype of mice carrying the human *SLC30A8* p.Arg138*

Since neither global nor tissue specific *Slc30a8* KD mouse models have recapitulated the human phenotype in carriers of the *SLC30A8* p.Arg138* variant, we tried to overcome this problem by using a mouse model carrying the *Slc30a8* p.Arg138* variant^11^. These mice do not express the truncated ZnT8 protein^11^. On a standard chow diet there was no evidence for enhanced insulin secretion^11^. However, we examined whether they might do so on a high fat diet (HFD). This was indeed the case (Fig. 6a-h), and the same differences in proinsulin/insulin and proinsulin/C-peptide ratios were seen as in humans. No changes were seen in insulin clearance.

**Fig. 6:**
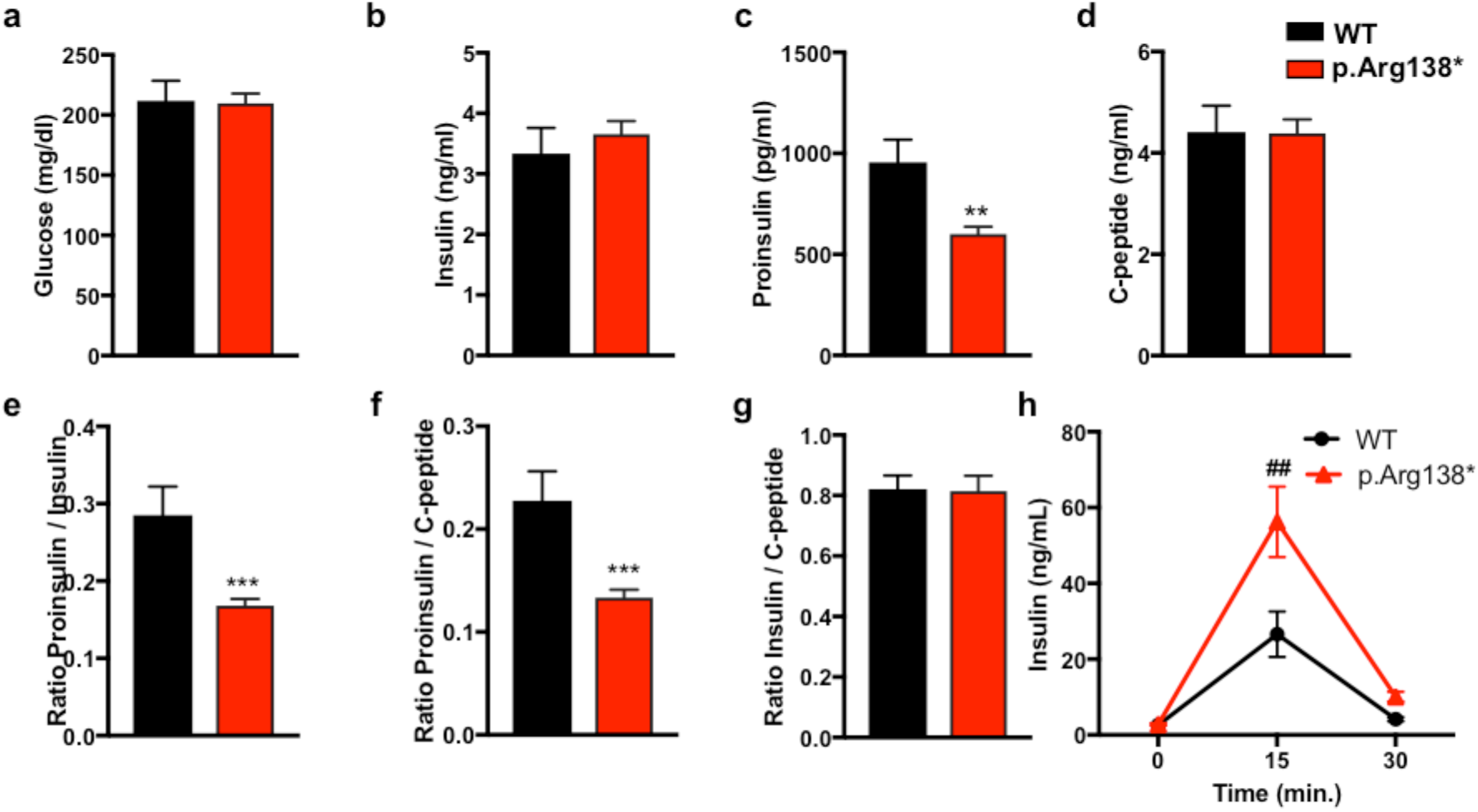
Male p.Arg138* mice on high-fat diet show enhanced insulin secretion and proinsulin processing. Circulating **a**, glucose **b**, insulin **c**, proinsulin **d**, C-peptide **e**, proinsulin/insulin ratio **f**, proinsulin/C-peptide ratio and **g**, insulin/C-peptide ratio in fasted WT and p.Arg138* mice (n= 10 WT, 17 p.Arg138*) after 20 weeks on HFD. **h**, Insulin response to oral glucose (2g/kg) exposure (n=5 WT, 11 p.Arg138*) after 30 weeks on HFD. p**<0.01, p***<0.005 using Students T test; p^##^<0.01 using two-way Anova.

### Impact of p.Arg138* on protein localization and cytosolic zinc distribution in INS-1 cells

Although we found no evidence in either mouse or our human β-cell model to support the presence of a truncated protein we explored the possibility of what might happen if a truncated protein resulted from mRNA evading NMD. Transient overexpression of tagged ZnT8-p.Arg138* fusion proteins in a rat insulinoma cell line, INS-1e, showed distinct punctate distribution patterns, consistent with localization of the truncated ZnT8 protein to secretory granules, as previously observed with the full length protein^20^ (Supplementary Fig. 5a-c) Additionally, Western blot showed stable expression of truncated ZnT8 in native INS1e cells (Supplementary Fig. 5d).

To investigate the effects of a truncated ZnT8 protein on cytosolic free Zn^2+^, we used a genetically-encoded Zn^2+^ sensor eCALWY-4^21^. Overexpression of the truncated protein (p.Arg138*) had no impact on cytosolic free Zn^2+^ when expressed in INS-1 WT cells ruling out a dominant negative effect for the truncated protein (Supplementary Fig. 5e-h).

### Influence of common *SLC30A8* variants p.Trp325Arg in primary human islets

While adult human islets show high levels of *SLC30A8* expression there was no reproducible effect of the p.Arg325Trp variant on *SLC30A8* expression in human islets from cadaveric donors (Fig. 7a). Islets obtained from cadaveric p.Trp325 carriers secreted more insulin than p.Arg325Arg carriers (Fig. 7b-e). The increased glucose responsiveness was observed at submaximal glucose stimulation (6 mM) rather than at maximal glucose stimulation (16.7 mM) (Fig. 7b-c). Increasing glucose from 1 mM to 6 mM stimulated insulin secretion 2.2- and 2.7-fold in p.Arg325 and p.Trp325 carriers respectively, with no effect on insulin content (Fig. 7c-d). This secretion pattern echoes the one observed after siRNA of *SLC30A8* KD in EndoC-βH1. Insulin secretion in p.Trp325 carriers was also increased at high glucose (16.7 mM) when co-exposed to depolarizing [K^+^]_o_ (70 mM) (Fig. 7e) as also seen after *SLC30A8* KD in EndoC-βH1.

**Fig. 7:**
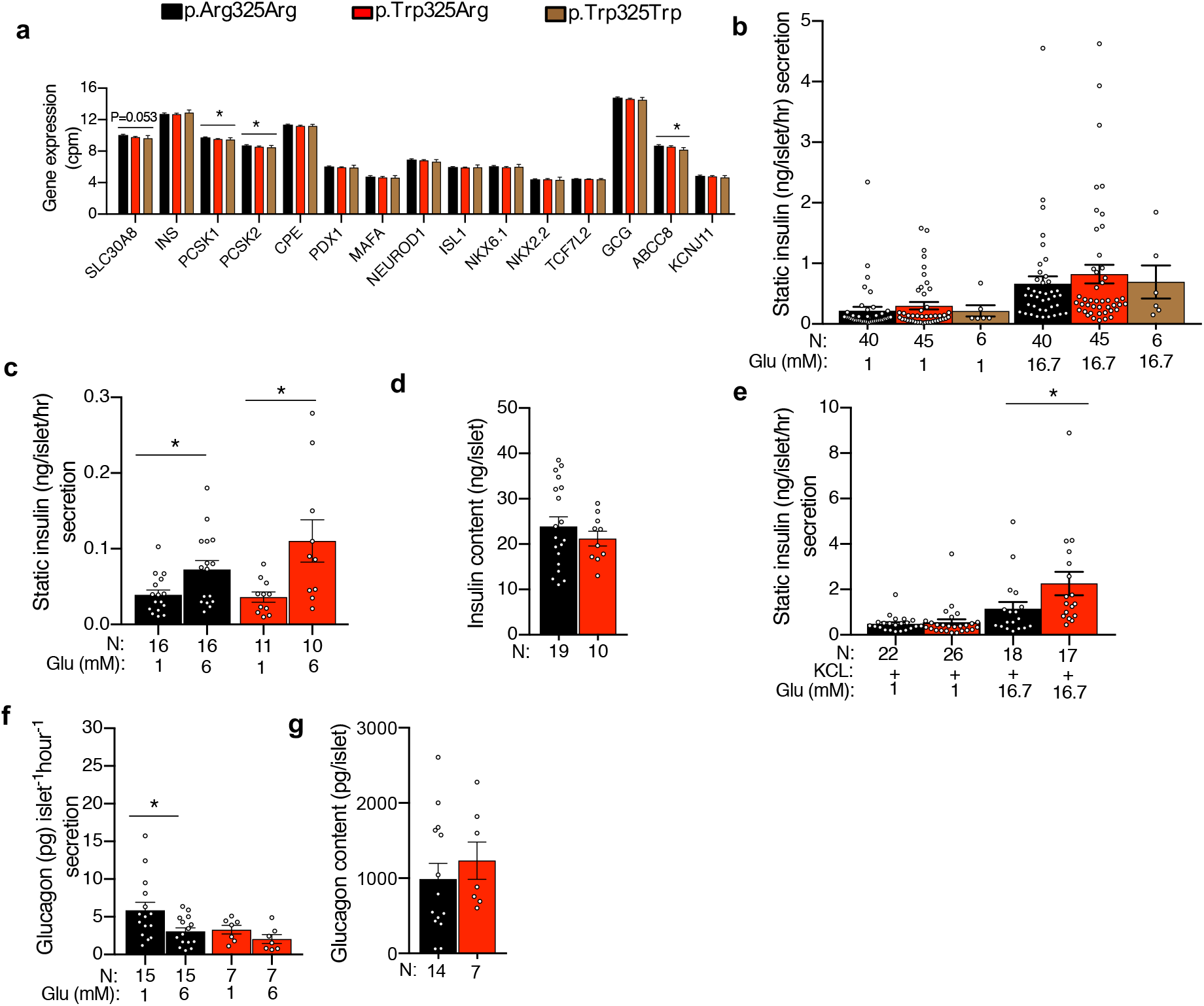
*SLC30A8*- p.Trp325 leads to enhanced insulin secretion in human islets. **a**, Effect of p.Trp325Arg genotype (p.Arg325Arg=66, p.Trp325Arg=63 and p.Trp325Trp=11) on expression of *SLC30A8* and other genes involved in insulin production, secretion and processing. **b**, Effect of p.Trp325Arg genotype on static insulin secretion in presence of low and high glucose stimulatory conditions. **c-d**, Effect of p.Trp325Arg genotype on static insulin secretion in (**c**) low stimulatory conditions and their (**d**) insulin contents. **e**, Effect of p.Trp325Arg genotype on static insulin secretion in presence of low and high glucose and KCL. **f**, Static glucagon response to glucose and **g**, glucagon content at basal glucose. Data are Mean ±SEM; Glu-glucose. Analysis by linear regression or Mann-Whitney test (Methods); * p<0.05.

As *SLC30A8* is highly expressed in human alpha cells^1^, we also measured glucagon secretion from the same islets (Fig. 7f). In islets from p.Arg325Arg donors, 6 mM glucose inhibited glucagon secretion by ~50% compared to 1 mM glucose. In islets from p.Trp325Arg donors, glucagon secretion at 1 mM glucose was reduced by 50% compared to p.Arg325Arg donors with no effect on glucagon content (Fig. 7f-g).

We also explored whether the p.Trp325Arg variant would have trans-eQTL effects on genes involved in insulin production and secretion^22^ (Fig. 7a). Expression of *PCSK1* (P=0.041) and *PCSK2* (P=0.045) were reduced. Among the genes encoding for K_ATP_ channels subunits only *ABCC8* (P=0.049) expression was significantly affected in islets from p.Trp325 carriers compared to non-carriers (Fig. 7a). Taken together, the data suggest the common T2D-protective allele (p.Trp325) may improve the response to a glucose challenge by enhancing insulin secretion and possibly by reducing glucagon secretion in primary human islets.

## Discussion

The current study demonstrates the strengths of using human models for studying the consequences of LoF mutations in humans, particularly by demonstrating a stronger protective effect of p.Arg138* in individuals carrying the common risk p.Arg325 allele on the same haplotype. However, the minor p.Trp325 allele was also associated with protection against T2D albeit less pronounced. This emphasizes the importance of taking into account the genetic background of the human LoF carrier.

Whilst the data from all our sub-studies are consistent with increased glucose responsiveness, the precise molecular mechanisms for these phenotypes, involvement of zinc and an explanation for why there are discrepancies between humans and rodents remain elusive. In the IPS-derived beta-like cells, the p.Arg138* variant dramatically lowered expression with evidence of NMD resulting in haploinsufficiency. Similarly, in the mouse model we were unable to detect the truncated protein, but we could detect appreciable levels of RNA^11^.

The most reproducible finding in all sub-studies of p.Arg138* was enhanced glucose-stimulated insulin secretion accompanied by increased conversion of proinsulin to C-peptide and insulin. Carriers of p.Trp325 displayed a similar phenotype, which is in line with a previous study showing impaired proinsulin conversion in carriers of the risk p.Arg325 allele^23^. There could also be other potential explanations for this effect, as it has been suggested that it takes some time for insulin to mature and become biologically active^24, 25^. It is possible that the pronounced effects of the LoF mutation at 20 and 40 min of test meal could reflect such a mechanism.

The present and previous studies demonstrate that loss of ZnT8 function after silencing the murine gene reduces total cellular zinc content as well as free Zn^2+^ in the cytosol and granules^7,10, 20, 26^. LoF p.Arg138* (assuming no or minimal escape from NMD) is therefore likely to exert the same effects on intracellular zinc concentrations and may thus impact insulin secretion through intracellular mechanisms, including potential differences in Zn^2+^ secretion. Also, a recent study showed that the p.Arg325Arg variant was associated with higher islet zinc concentrations^27^. In the present study over-expression of the LoF mutation p.Arg138* in INS-1 cells did not result in changes in cytosolic zinc concentrations leaving a reduction of zinc in insulin granules as a plausible explanation which still needs to be experimentally confirmed.

In support of a protective effect of lowering intracellular zinc concentrations on development of diabetes, in the CNS, Zn^2+^ plays an important role as a regulator of cellular excitability^28^ and Zn^2+^ has been reported to activate K_ATP_ channels^29^, inhibit L-type voltage-gated Ca^2+^ channels and inhibit insulin secretion^30^.

Taken together, our data consistently demonstrate that heterozygosity for a LoF mutation p.Arg138* and homozygosity for a common variant p.Trp325Trp of the *SLC30A8* are associated with increased insulin secretion capacity and lower risk of T2D without any negative effect. Therefore, ZnT8 remains an appealing safe target for antidiabetic therapy preserving β-cell function.

## Methods

### Human study population

The Botnia Study has been recruiting patients with T2D and their family members in the area of five primary health care centers in western Finland since 1990. Individuals without diabetes at baseline (relatives or spouses of patients with T2D) have been invited for follow-up examinations every 3-5 years^12^. The Prevalence, Prediction and Prevention of diabetes (PPP)–Botnia Study is a population-based study in the same region including a random sample of 5,208 individuals aged 18 to 75 years from the population registry^31^. Diabetes Registry Vaasa (DIREVA) is regional diabetes registry of > 5000 diabetic patients from Western Finland (Botnia region)^32^. In the current study, we included >14,000 individuals (Botnia family study=5678, PPP=4862, and DIREVA=3835). All participants gave their written informed consent and the study protocol was approved by the Ethics Committee of Helsinki University Hospital, Finland (the Botnia studies) and the Ethics Committee of Turku University Hospital (DIREVA).

#### Oral Glucose Tolerance Test (OGTT) and test meal

Subjects maintained a weight-maintaining diet and avoided vigorous exercise for 3 days prior to the OGTT or test meal, which were performed after an overnight fast. Height, weight, hip and waist circumferences, fat percentage (%, bioimpedance analyzer) and blood pressure (sitting, 3 measurements after 5 min rest) were measured. The participants ingested 75 g dextrose (in a couple of minutes, OGTT) or a 526 kcal mixed meal (in 10 minutes, test-meal: 76 g carbohydrate, 17 g protein and 15 g fat). Blood samples were drawn from an antecubital vein for plasma (P-) glucose and serum (S-) insulin and C-peptide at 0, 30, 120 min during the OGTT; for P-glucose, P-glucagon, S-insulin, S-C-peptide, S-zinc, and total S-GLP-1 at 0, 20, 40, 70, 100, 130, 160 and 190 min during the test meal. Test meal samples for S-FFA were collected at 0, 40 and 120 min and for S-proinsulin at 0, 20, 40 and 130 min, respectively. Urine was collected between 0 – 70 and 70 – 190 min for the determination of glucose and zinc excretion during the test meal.

#### Intravenous Glucose Tolerance Test (IVGTT)

IVGTT group consists of total 849 (male-403, female-446) individuals with an average age of 51 years. An antecubital polyethylene catheter was placed to one hand for the infusion of 0.3 g/kg body weight of glucose (maximum dose 35 g) intravenously for 2 min. A retrogradely positioned wrist vein catheter was placed in the other hand, held in a heated (70°C) box in order to arterialize the venous blood. Arterialized blood samples were drawn at 0, 2, 4, 6, 8,10, 20, 30, 40, 50 and 60 min for P-glucose and S-insulin.

#### Biochemical measurements

P-glucose was analyzed using glucose oxidase (Beckman Glucose Analyzer, Beckman Instruments, Fullerton, CA, USA; Botnia Family Study) or glucose dehydrogenase method (Hemocue, Angelholm, Sweden; PPP-Botnia and test meal studies). In the Botnia Family study, S-insulin was measured by radioimmunoassay (RIA, Linco; Pharmacia, Uppsala, Sweden), enzyme immunoassay (EIA; DAKO, Cambridgeshire, U.K.) or fluoroimmunometric assay (FIA, AutoDelfia; Perkin Elmer Finland, Turku, Finland). For the analysis, insulin concentrations obtained with different assays were transformed to cohere with those obtained using the EIA. The correlation coefficient between RIA and EIA as well as between FIA and EIA was 0.98 (P < 0.0001). S-insulin was measured by the FIA in baseline visit of PPP-Botnia and the test meal study (correlation co-efficient 0.98). S-proinsulin was measured using RIA (Linco; Pharmacia, Uppsala, Sweden, OGTT data) or EIA (Mercodia AB, Uppsala, Sweden; test-meal data), and P-glucagon using RIA (EMD Millipore, St. Charles, MO; OGTT data) or EIA (Mercodia AB, Uppsala, Sweden; test-meal data). S-FFA was measured by an enzymatic colorimetric method (Wako Chemicals, Neuss, Germany). Serum total cholesterol, HDL and triglyceride concentrations were measured with Cobas Mira analyzer (Hoffman LaRoche, Basel, Switzerland), and since 2006 with an enzymatic method (Konelab 60i analyser; Thermo Electron Oy, Vantaa, Finland). Serum LDL cholesterol was calculated using the Friedewald formula. Blood collected in tubes containing DPP4-inhibitors was used for radioimmunoassay^33^ for total P-GLP-1 (intact GLP-1 and the metabolite GLP-1 9-36 amide) during test meal.

Serum and urine samples for zinc were collected in trace element tubes (Beckton Dickinson, NJ, USA) and S- and U-zinc analyzed by two commercial laboratories: NordLab (Oulu, Finland; atom absorption spectrophotometry, AAS) until 6^th^ May 2015, then in Synlab (Helsinki, Finland; AAS for serum, mass spetrophotometry ICP-MS for U-zinc). The S-zinc concentrations were corrected for P-albumin (r = 0.34, p 0.008 Nordlab, r = 0.34, p 0.03 Synlab).

Corrected insulin response (CIR) was calculated for test meal (at 20 min) and OGTT (at 30 min.) using the formula CIR(t)= Ins(t) /[Gluc(t) • (Gluc(t)-3.89)], where Ins(t) and Gluc(t) are insulin (in mU/L) and glucose concentrations (in mmol/L) at sample time point t (min)^34^. Estimation of Insulin clearance index was done on the model based estimation of glucose-, insulin- and C-peptide curves during the test meal using the equation AUC(ISR) / [(AUC(ins)+(I(basal)-I(final) • MRT(ins)], where AUC(ISR) is the area under the curve of insulin secretion rate, AUC(ins) is the area under the curve of insulin concentration, I(final) is insulin concentration at the end, and I(basal) insulin concentration at the beginning of the study^35^. MRT(ins) is the mean residence time of insulin, and was assumed to be 27 minutes as reported previously^36^.

#### Genotyping

We analyzed genotype data for rs13266634 (p.Trp325Arg) and rs200185429 (p.Arg138*) for three cohorts genotyped with different genome-/exome-wide chips: the Botnia family cohort (Illumina Global Screening array-24v1, genotyped at Regeneron Pharmaceuticals), PPP-Botnia (Illumina HumanExome v1.1 array, genotyped at Broad Institute^3^) and DIREVA (Illumina Human CoreExome array-24v1, genotyped at LUDC). For the Botnia family cohort, genotype data for p.Arg138* were imputed (info score >0.95) from the available GWAS data by phasing using SHAPT-IT v2^37^ and imputing using the GoT2D reference panel^38^ by IMPUTEv2^39^ . The carrier status of imputed p.Arg138* was additionally confirmed from exome sequencing data. Genotyping (p.Trp325Arg and p.Arg138*) the family members participating in the genotype based recall study (test meal study) was performed using TaqMan (Applied Biosystems, Carlsbad, CA). The genotype distribution of both variants was in accordance with Hardy-Weinberg equilibrium in all the cohorts. We did not detect any Mendelian errors in the families.

#### Genetic Association Analysis

All the quantitative traits were inversely normally transformed before the analyses. The family-based recall study included only non-diabetic subjects during test meal and analysis of data was performed using family-based association analyses adjusting for age, sex, BMI, and other covariates if appropriate, using QTDT (v2.6.1)^40^. The significance levels were derived from 100,000 permutations as implemented in QTDT. Also, the OGTT study included only non-diabetic subjects. The association analysis was performed using mixed linear model considering genetic relatedness among samples as implemented in GCTA (v1.91)^41^.

#### Study participants and their clinical measurements in Verona Newly Diagnosed Diabetes Study (VNDS)

The Verona Newly Diagnosed Type 2 Diabetes Study (VNDS; NCT01526720) is an ongoing study aiming at building a biobank of patients with newly diagnosed (within the last six months) type 2 diabetes. Patients are drug-naïve or, if already treated with antidiabetic drugs, undergo a treatment washout of at least one week before metabolic tests are performed^42^. Each subject gave informed written consent before participating in the research, which was approved by the Human Investigation Committee of the Verona City Hospital. Metabolic tests were carried out on two separate days in random order^42^. Plasma glucose concentration was measured in duplicate with a Beckman Glucose Analyzer II (Beckman Instruments, Fullerton, CA, USA) or an YSI 2300 Stat Plus Glucose&Lactate Analyzer (YSI Inc., Yellow Springs, OH, USA) at bedside. Serum C-peptide and insulin concentrations were measured by chemiluminescence as previously described^42^. The analysis of the glucose and C-peptide curves during the OGTT was carried out with a mathematical model as described previously^42^. This model was implemented in the SAAM 1.2 software (SAAM Institute, Seattle, WA) to estimate its unknown parameters. Numerical values of the unknown parameters were estimated by using nonlinear least squares. Weights were chosen optimally, i.e., equal to the inverse of the variance of the measurement errors, which were assumed to be additive, uncorrelated, with zero mean, and a coefficient of variation (CV) of 6-8%, A good fit of the model to data was obtained in all cases and unknown parameters were estimated with good precision. In this paper we report the response of the beta cell to glucose concentration (proportional control of beta cell function), which in these patients accounts for 93.2±0.3% of the insulin secreted by the beta cell in response to the oral glucose load. Genotypes were assessed by the high-throughput genotyping Veracode technique (Illumina Inc, CA), applying the GoldenGate Genotyping Assay according to manufacturer’s instructions. Hardy-Weinberg equilibrium was tested by chi-square test. Variant association analyses were carried out by generalized linear models (GLM) as implemented in SPSS 25.0 and they were adjusted for a number of potential confounders, including age, sex and BMI.

### iPSC generation, differentiation and genome editing

#### iPSC generation and maintenance

The human induced pluripotent stem cell line (hiPSC) SB Ad3.1 was previously generated and obtained through the IMI/EU sponsored StemBANCC consortium via the Human Biomaterials Resource Centre, University of Birmingham (http://www.birmingham.ac.uk/facilities/hbrc). Human skin fibroblasts were obtained from a commercial source (Lonza CC-2511, tissue acquisition number 23447). They had been collected from a Caucasian donor with no reported diabetes with fully informed consent and with ethical approval from the National Research Ethics Service South Central Hampshire research ethics committee (REC 13/SC/0179). The fibroblasts were reprogrammed to pluripotency as previously described^43^ and were subjected to the following quality control checks: SNP-array testing via Human CytoSNP-12v2.1 beadchip (Illumina #WG-320-2101), DAPI-stained metaphase counting and mFISH, flow cytometry for pluripotency markers (BD Biosciences #560589 and 560126), and mycoplasma testing (Lonza #LT07-118).

#### CRISPR-Cas9 mediated generation of p.Arg138* human induced pluripotent stem cell line

Several guide RNAs (gRNAs) were designed using MIT CRISPR tool (http://crispr.mit.edu/) to target near exon 3 of *SLC30A8* (ENST00000456015). The gRNAs were also subjected to an additional BlastN search (www.ensembl.org) to confirm specificity and identified no additional off-target sites. The gRNA (AGCAGGTACGGTTCATAGAG) was sub-cloned into the *BsbI* restriction sites in pX330^44^ plasmid that was previously modified to contain a puromycin selection cassette. Single strand oligonucleotides for homology-directed repair (HDR) were synthesised by Eurogentec, stabilised by addition of a phosphorothioate linkage at the 5’ end, and contained two nucleotide changes: i) the T2D-protective nonsense mutation at codon-138 (c.412C>T, p.Arg138*), which also mutated the PAM sequence, and ii) a silent missense mutation at codon-139 (c.417A>T, p.Ala139Ala) to introduce an *Alu*I restriction site for genotyping.

Human iPSCs were co-transfected with *SLC30A8*-px330-puromycin resistant vectors and HDR oligos using Fugene6 according to manufacturer’s guidelines (Promega #E2691). Following transient puromycin-selection, single clones were picked and expanded as described previously^45^. Genotyping PCR was performed using primers (Forward: TACCCCAGGAATGGCTTCTC; Reverse: ACGTGTTCCTGTTGTCCCA) to amplify targeted region followed by *Alu*I restriction digest. Successfully targeted clones were confirmed via Sanger sequence and monoallelic sequencing was performed by TA-cloning (pGEM^®^-T Easy Vector System; Promega #A1360) of the PCR amplicons. The control hiPSC line (p.Arg138Arg) was generated from hiPSC cells that went through the CRISPR pipeline without being edited at the *SLC30A8* locus. The two p.Arg138* clones (A3 and B1) and the unedited control line (p.Arg138Arg) passed quality control checks that included repeat chromosome counting and pluripotency testing.

#### In vitro differentiation of hiPSCs towards Beta-like cells

Directed differentiation of hiPSCs towards beta-like cells was performed using a previously published protocol^14, 46^. hiPSCs were seeded on Growth Factor Reduced Matrigel-coated CellBind 12-well tissue culture plates (Corning #356230 & #3336) at a cell density of 1.3×10^6^ in mTesR1 (Stem Cell Technologies #05850) with 10 μM Y-27632 dihydrochloride (Abcam #ab120129). The following morning, hiPSCs were fed mTesR1 media >4 hours before starting the seven-stage differentiation protocol.

##### Stage 1 (Definitive Endoderm)

Cells were washed once with PBS before adding 0.5% bovine serum albumin (BSA; Roche #10775835001) MCDB131 media [(ThermoFisher Scientific #10372019) containing 1x Penicillin-Streptomycin (Sigma #P0781), 1.5 g/L sodium bicarbonate (ThermoFisher Scientific #25080060), 1x GlutaMAX™(ThermoFisher Scientific #35050038) and 10 mM Glucose (ThermoFisher Scientific #A2494001)] supplemented with 100 ng/mL Activin A (Peprotech #120-14) and 3 μM CHIR 99021 (Axon Medchem #1386). On day 2 and 3, cells were cultured with 0.5% BSA MCDB131 media supplemented with either 100 ng/mL Activin A and 0.3 μM CHIR 99021 (day 2) or with 100 ng/mL Activin A alone (day 3).

##### Stage 2 (Primitive Gut Tube)

Cells were cultured for 48 hours in 0.5% BSA MCDB131 media with 0.25 mM ascorbic acid (Sigma #A4544) and 50 ng/mL KGF (PeproTech #100-19).

##### Stage 3 (Posterior Foregut)

Cells were cultured for two days in 2% BSA MCDB131 media supplemented with 1 g/L sodium bicarbonate, 0.25 mM ascorbic acid, 0.5x Insulin-Transferrin-Selenium-Ethanolamine (ITS-X; ThermoFisher Scientific #51500056), 1 μM retinoic acid (RA; Sigma-Aldrich #R2625), 0.25 μM Sant-1 (Sigma-Aldrich #S4572), 50 ng/ml KGF, 100 nM LDN193189 (Stemgent #04-0074), and 100 nM α-Amyloid Precursor Protein Modulator (Merck #565740).

##### Stage 4 (Pancreatic Endoderm)

Cells were cultured for three days in 2% BSA MCDB131 media supplemented with 1 g/L sodium bicarbonate, 0.25 mM ascorbic acid, 0.5x ITS-X, 0.1 μM RA, 0.25 μM Sant-1, 2 ng/ml KGF, 200 nM LDN193189 and 100 nM α-Amyloid Precursor Protein Modulator.

##### Stage 5 (Endocrine Progenitors)

Cells remained in planar culture for three days in 2% BSA MCDB131 media supplemented with 20 mM final glucose, 0.5x ITS-X, 0.05 μM RA, 0.25 μM Sant-1, 100 nM LDN193189, 10 μM ALK5 Inhibitor II (Enzo Life Sciences #ALX-270-445), 1 μM 3,3,5-Triiodo-L-thyronine sodium salt (T3; Sigma-Aldrich #T6397), 10 μM zinc sulfate heptahydrate (Sigma # Z0251), and 10 μg/mL heparin sodium salt (Sigma #H3149).

##### Stage 6 (Endocrine Cells)

Cells remained in planar culture for six days in 2% BSA MCDB131 media supplemented with 20 mM final glucose, 0.5x ITS-X, 100 nM LDN193189, 10 μM ALK5 Inhibitor II, 1 μM T3, 10 μM zinc sulfate heptahydrate, and 100 nM γ-Secretase Inhibitor XX (Merck Millipore #565789).

##### Stage 7 (Beta-like Cells)

Cells remained in planar culture for another six days in 2% BSA MCDB131 media supplemented with 20 mM final glucose, 0.5x ITS-X, 10 μM ALK5 Inhibitor II, 1 μM T3, 1 mM N-Cys (Sigma-Aldrich #A9165), 10 μM Trolox (EMD Millipore #648471), 2 μM R248 (SelleckChem #S2841), and 10 μM zinc sulfate heptahydrate.

#### Quantification of SLC30A8 gene expression in Beta-like Cells derived from CRISPR-edited hiPSCs

Expression of *SLC30A8* was measured at the end of stage 7 using quantitative PCR (qPCR). Briefly, RNA was extracted using TRIzol Reagent (Life Technologies #15596026) according to manufacturer’s instructions. cDNA was amplified using the GoScript Reverse Transcription Kit (Promega #A5000). qPCR was performed using 40 ng of cDNA, TaqMan^®^ Gene Expression Master Mix (Applied Biosystems #4369017) and primer/probes for *SLC30A8* (Hs00545182_m1) or the housekeeping gene *TBP* (Hs00427620_m1). Gene expression was determined using the ∆∆CT method by first normalizing to *TBP* and then to the control p.Arg138Arg sample (n=6-7 wells from two differentiations).

#### Allele-specific SLC30A8 expression in Beta-like Cells derived from CRISPR-edited hiPSCs

Stage 7 cells were treated with 100 μg/mL cycloheximide (Sigma #C4859) or DMSO (Sigma #D2650) for four hours at 37°C^47^ before harvesting for RNA and cDNA synthesis as above. Allele specific expression was measured using the QX10 Droplet Digital PCR System and C1000 Touch Thermal Cycler according to manufacturer’s guidelines (Bio-Rad). Custom primers and probes for the detection of p.Arg138* variant were designed using Primer3Plus (Applied Biosystems): Forward primer AGTCTCTTCTCCCTGTGGTT; Reverse primer ATGATCATCACAGTCGCCTG; FAM probe 5’-FAM-ATGGCACCGAGCTGA-MGB-3’; VIC probe 5’-VIC-ATGGCACTGAGCTGAGA-MGB-3’. Results were analysed using Quanta Soft software (Bio-Rad) and presented as a ratio of wildtype to HDR-edited allele expression (n>3 wells from two differentiations).

## EndoC-βH1 culture

The results obtained in EndoC-βH1 are from two distinct teams (Helsinki and Oxford) with different batches of EndoC-βH1 cultures. Here, we report both methods and specify for each experiment the origin of the culture (Helsinki or Oxford). EndoC-βH1 cells were cultured in medium and grown on a matrix as described previously^48^ and tested negative for mycoplasma.

### SLC30A8 knockdown in EndoC-βH1 cells

In Oxford, EndoC-βH1 cells were transfected with 10 nM siRNA (either SMARTpool ON-TARGETplus SLC30A8 or scramble [Dharmacon #L-007529-01]) and Lipofectamine RNAiMAX (Life Technologies #13778-075) according to manufacturer’s instructions for a total of 72 hours. In Helsinki, EndoC-βH1cells were transfected using Lipofectamin RNAiMAX (life technologies). 20nM siRNA ON-TARGET*plus* siRNA SMARTpool for human *SLC30A8* gene (Dharmacon; L-007529-01) and ON-TARGET*plus* Non-targeting pool (siNT or Scramble) (Dharmacon; D-001810-10-05) were used following the protocol as described previously^49^. Cells were harvested 96 h post-transfection for further studies.

### Insulin secretion measurements in EndoC-βH1 cells

In Oxford, cells were subjected to static insulin secretion assays 72hrs after siRNA transfection as described previously^50^, apart from the following modifications: cells were stimulated for 1 hr with 1 mM glucose, 20 mM glucose, 1 mM glucose + 200 μM tolbutamide, or 20 mM glucose + 500 μM diazoxide. Insulin levels were measured in both supernatants and cells using the Insulin (human) AlphaLISA Detection Kit and EnSpire Alpha Plate Reader (Perkin Elmer #AL204C and #2390-0000, respectively). Cell count per well was measured via CyQUANT Direct Cell Proliferation Assay (Thermo Fisher# C35011). Data are presented as insulin secretion normalized to percentage of insulin content from Control condition. RNA extraction, cDNA synthesis, and qRT-PCR was performed as above (*SLC30A8* gene expression in CRISPR-edited hiPSCs derived beta like cell section) to determine *SLC30A8* knockdown and expression of the K_ATP_ channel genes (*ABCC8* Hs01093752_m1 and *KCNJ11* Hs00265026_s1; ThermoFisher Scientific). In Helsinki, EndoC-βH1 cells were transfected with 20nM siRNA and Scramble control. Following 96h of siRNA transfection, cells were incubated overnight in 1 mM glucose containing EndoC-βH1 culture medium. One hour prior to glucose stimulation assay, the media was replaced by βKREBS (Univercell Biosolution S.A.S., France) without glucose. Cells were stimulated with 16.7 mM glucose and 50 mM KCl (Sigma-Aldrich) in βKREBS for 30 min at 37°C in a CO_2_ incubator. The cells were then washed and lysed with TETG (Tris pH8, Trito X-100, Glycerol, NaCl and EGTA) solution (Univercell Biosolution S.A.S., France) for the measurement of total insulin content. Secreted and intracellular insulin were measured using a commercial human insulin Elisa kit (Mercodia AB, Uppsala, Sweden) as per manufacturer’s instructions (Helsinki).

### Electrophysiological measurements in EndoC-βH1 cells (Oxford)

*SLC30A8* was knocked down in EndoC-βH1 as above. K^+^_ATP_ channel conductance was measured in a perforated patch whole cell configuration, and patch-clamped using an EPC 10 amplifier and HEKA pulse software. KREBS extracellular solution was perfused in at 32°C and contained: 138 mM NaCl, 3.6 mM KCl, 0.5 mM MgSO_4_, 10 mM HEPES, 0.5 mM NaH_2_PO_4_, 5 mM NaHCO_3_, 1.5 mM CaCl_2_, 1 mM glucose and 100 μM Diazoxide (Sigma-Aldrich #D9035). The perforation of the membrane was achieved using an intra-pipette solution containing: 0.24 mg/mL amphotericin B, 128 mM K-gluconate (Sigma #Y0000005 and G4500 respectively), 10 mM KCl, 10 mM NaCl, 1 mM MgCl_2_, 10 mM HEPES, pH 7.35 (KOH). Conductance data are normalised to cell size and presented as pS.pF^-1^. Expression of *ABCC8, KCNJ11, B2M*, and *TBP* were measured via qPCR as above (*SLC30A8* gene expression in CRISPR-edited hiPSCs derived beta like cell section).

### Insulin and Proinsulin secretion and content (Helsinki)

For the measurement of secreted insulin or proinsulin in the supernatant, 96h post-transfected cells were washed twice with 1X PBS and incubated with fresh EndoC-βH1 culture medium for next 24h. Secreted and intracellular insulin and proinsulin were measured using a commercial human insulin Elisa and human proinsulin Elisa kit from Mercodia (Mercodia AB, Uppsala, Sweden). Total cellular protein content was also determined with the BCA protein assay kit (Thermo Scientific, Pierce). Proinsulin to insulin ratio was calculated by dividing the respective values measured from the supernatant and the cells (pmol/L).

### Immunoblotting (Helsinki)

Total cellular protein was prepared with Laemmli buffer and resolved using Any kD Mini-Protean-TGX gel (Bio-Rad). Immunoblot analysis was performed by overnight incubation of with primary antibodies against ZNT8 (Abcam; #ab136990; 1:500), PC1/3 (Cell Signaling; #11914; 1:1000), CPE (BD Bioscience; #610758; 1:1000), Phospho-AKT-Ser473 (Cell Signaling; #4060; 1:1000), AKT (Santa-Cruz; #SC-8312; 1:500). The membranes were further incubated with species-specific HRP-linked secondary antibodies (1:5000) and visualization was performed following ECL exposure with ChemiDoc XRS+ system and Image Lab Software (Bio-Rad). A loading control of either alpha-Tubulin (Sigma; T5168; 1:5000) or beta-actin (Sigma; A5441; 1:5000) was performed on the same blot for all western blot data. Densitometric analysis of bands from image were calculated using Image J (Media Cybernetics) software and intensities compared as ZNT8, PC1/3, phosphor-AKT-Ser473 to tubulin; CPE to beta-actin.

### Cell viability assay, MTT (Helsinki)

EndoC-βH1 cells were transfected with either siScramble or siSLC30A8 for 96h. The viability of cells after 24 h of tunicamycin (10 μg/ml) treatment was determined using Vybrant MTT Cell proliferation kit (ThermoFisher Scientific; #M6494), the standard MTT [3-(4,5-dimethylthiazol-2-yl)-2,5-diphenyltetrazolium bromide] assay. All the treatments were performed on cells with equal seeding density (5×10^4^ cells/well) in 96 wells plate. The purple formazan crystals generated after 2 h incubation with MTT buffer were dissolved in DMSO, and the absorbance was recorded on a microplate reader at a wavelength of 540nm.

### RNA (mRNAs) sequencing of EndoC-βH1 cells

For RNA sequencing post 96h siScramble (n=3) or siSLC30A8 (n=3) transfected EndoC-βH1 cells were used and the total RNA was extracted with Macherey-Nagel RNA isolation kit as per manufacturer’s instruction. RNA sequencing was performed using Illumina TruSeq-mRNA library on NextSeq 500 system (Illumina) with an average of >15 million paired-end reads (2 × 75 base pairs). RNA sequencing reads were aligned to hg38 using STAR (Spliced Transcripts Alignment to Reference)^51^, genome annotations were obtained from the GENCODE (Encyclopedia of Genes and Gene Variants) v22^52^ program, and reads counting were done using featureCounts^53^. Further downstream analysis was perform using edgR^54^ software package, low expressed (<1 average count per million) genes were removed, read counts were normalized using TMM^55^ (trimmed mean of M-values), differential expression analysis was performed using method similar to Fisher’s Exact Test and corrected for multiple testing using FDR (1%).

### Data Analyses

Data are reported as mean (SEM). Statistical analyses were performed using Prism 6.0 (GraphPad Software). All parameters were analyzed using Mann-Whitney test or Unpaired Student’s t-test as indicated.

## Mouse Model

### Animals

All procedures were conducted in compliance with protocols approved by the Regeneron Pharmaceuticals Institutional Animal Care and Use Committee. The *Slc30a8*^Tgp.Arg138*^mouse line is made in pure C57Bl/6 background by changing nucleotide 409 from T into C in exon 3, which changes the arginine into a stop codon^11^. The mutated allele has a self-deleting neomycin selection cassette flanked by loxP sites inserted at intron 3, deleting 29 bp of endogenous intron 3 sequence. Mice were housed (up to five mice per cage) in a controlled environment (12-h light/dark cycle, 22C, 60–70% humidity) and fed *ad libitum* with either chow (Purina Laboratory 23 Rodent Diet 5001, LabDiet) or high-fat diet (Research Diets, D12492; 60% fat by calories) starting at age of 20 weeks. All data shown are compared to their respective WT littermates.

### Glucose Tolerance Test

Mice were fasted overnight (16 hr) followed by oral gavage of glucose (Sigma) at 2 g/kg body weight. Blood samples were obtained from the tail vein at the indicated times and glucose levels were measured using the AlphaTrak2 glucometer (Abbott). Submandibular bleeds for insulin were done at 0, 15, and 30 min post-injection.

### Hormone measurements

Submandibular bleeds of either overnight fasted or fed animals were done in the morning. Plasma insulin or proinsulin was analyzed with the mouse insulin/proinsulin EIA (Mercodia AB, Uppsala, Sweden), and C-peptide with the mouse C-peptide EIA (ALPCO). All EIAs were performed according to the manufacturer’s instructions.

### Data Analyses for mouse studies

Data are reported as mean (SEM). Statistical analyses were performed using Prism 6.0 (GraphPad Software). All parameters were analyzed by two-way ANOVA or Unpaired Student’s t-test as indicated.

## Expression of p.Arg138* mutation in INS1E

INS-1E cells^56^ were used for transient transfection of pcDNA3.1(+)-p.Arg138* construct fused to fluorescent m-Cherry at C-terminus using transfection reagent Viromer according to the manufacturer’s instructions. After transfections cells were collected at 24, 48, 72 and 96 hours and analysed by western blot analysis using mCherry (600-401-P16, Rockland) antibody. Untransfected cells were used as control and tubulin as a loading control. Two days after transient transfections with either p.Arg138*-mCherry (INS1E), p.Arg138*-HA or p.Arg138*-Myc-His construct (INS1E), cells were washed with PBS twice and fixed using 4% paraformaldehyde for 15 min at room temperature. Cells were permeabilized with 0.2 % Triton X-100 in phosphate-buffered saline (PBS) for 10 mins and to prevent unspecific binding were further blocked for 1 h with 5% FBS in PBS. INS1E cells transfected with either p.Arg138*-HA or p.Arg138*-Myc-His construct were incubated with the primary antibody (HA antibody: MMS-101P, Biolegend; His antibody: D291-A48, MBL; insulin antibody: A0564, DAKO), overnight at 4°C. Secondary antibodies were conjugated to Alexa Fluor 488 (Molecular Probes). Cells transfected with mCherry construct were imaged after 48 and 96 hours (INS1E) in order to visualize subcellular localization at different time points.

## Measurements of cytosolic zinc in INS-1(832/13) cells

### Cell culture

INS-1 (823/13) cells were grown in RPMI 1640 medium (Sigma-Aldrich, UK) supplemented with 10% (v/v) foetal bovine serum (FBS), 2 Mm L-glutamine, 0.05 mM 2-mercaptoethanol, 10 mM HEPES (Sigma-Aldrich), 1 mM sodium pyruvate (GIBCO, France), 2 mM L-glutamine and antibiotics (100 μg/ml Streptomycin and 100 U/ml penicillin). Cells were maintained in 95% oxygen, 5% carbon dioxide at 37°C.

### Co-transfection

Cells were seeded on sterile coverslips at 60% confluence and co-transfected using lipofectamine 2000 (Invitrogen, USA) according to the manufacturer’s instructions, with either the empty construct (EV) or the rare-truncated variant (c-Myc tag, R138X) construct and the Förster Resonance Energy transfer sensors (FRET), eCALWY-4 vector (free cytosolic zinc measurements).

### Protein extraction and Western (immuno-) blotting analysis

For protein extraction, RIPA buffer (1% Triton X-100, 1% sodium deoxycholate, 0.1% SDS, 0.15 mM NaCl, 0.01 M sodium Phosphate pH7.2) was used for lysis. Protein extracts were resolved in SDS-page (12% vol/vol acrylamide) and transferred to a polyvinylidene fluoride (PVDF) membrane, followed by blocking for 1 hour, immunoblotting with either c-Myc anti-mouse SLC30A8 (1:400) and the secondary anti-mouse antibody (1:10000, Abcam), and then the mouse monoclonal anti-tubulin (1:10000) and secondary anti-mouse for tubulin (1:5000). Chemiluminescence detection reagent (GE Healthcare) was used before exposing to hyperfilms.

### Immunocytochemistry

Cells were fixed in 4% (v/v) Phosphate-buffered saline/Paraformaldehyde (PFA). Cells were permeabilized in 0.5% (w/v) PBS/TritonX-100 and further saturated with PBS/BSA 0.1%. Cells were then incubated for 1 hour with the primary antibody, anti-c-Myc mouse antibody (1:200) followed by the secondary Alexa Fluor^®^ 568 nm anti-mouse IgG (H+L, 1:1000 Life Technologies, USA). Coverslips were mounted with mounting medium containing DAPI (Vectashield, USA) on microscope slides (ThermoScientific). Imaging was performed on a Nikon Eclipse Ti microscope equipped with a 63x/1.4NA objective, spinning disk (CAIRN, UK) using a 405, 488 and 561 nm laser lines, and images were acquired with an ORCA-Flash 4.0 camera (Hamamatsu) Metamorph software (Molecular Device) was used for data capture.

### Cytosolic free Zn^2+^ measurements

Acquisitions were performed 24 hours after transfection using an Olympus IX-70 wide-field microscope with a 40x/1.35NA oil immersion objective and a zyla sCMOS camera (Andor Technology, Belfast, UK) controlled by Micromanager software. Excitation was provided at 433 nm using a monochromator (Polychrome IV, Till Photonics, Munich, Germany). Emitted light was split and filtered with a Dual-View beam splitter (Photometrics, Tucson, Az, USA) equipped with a 505dcxn dichroic mirror and two emission filters (Chroma Technology, Bellows Falls, VT, USA - D470/24 for cerulean and D535/30 for citrine). Cells were perfused for 4 minutes with KREBS buffer (140 mM NaCl, 3.6 mM KCl, 0.5 mM NaH_2_PO_4_, 0.2 mM MgSO_4_, 1.5 mM CaCl_2_, 10 mM HEPES, 25 mM NaHCO_3_) without additives, next the buffer was changed to KREBS buffer containing 50 *μ*M N,N,N’,N’-tetrakis(2-pyridylmethyl)ethylenediamine (TPEN, Sigma) for 5 minutes, followed by perifusion with KREBS buffer containing 100 *μ*M ZnCl_2_ and 5 *μ*M of the Zn^2+^-specific ionophore 2-mercaptopyridine N-oxide (Pyrithione, Sigma). Image analysis was performed using ImageJ software. Steady-state fluorescence intensity ratio of acceptor over donor was measured, followed by the determination of the minimum and maximum ratios to calculate the free Zn^2+^ concentration using the following formula: [Zn^2+^] = Kd ∙ ((R – Rmin)/(Rmax – R)), in which Rmin is the ratio in the Zn^2+^ depleted state, after addition of 50 *μ*M TPEN, and Rmax was obtained upon Zn^2+^ saturation with 100 *μ*M ZnCl_2_ in the presence of 5 *μ*M pyrithione.

## Human Pancreatic islets

Experiments on primary human pancreatic islets were independently performed in two places 1) Oxford and 2) Lund university diabetes center (LUDC)

### Human pancreatic islets from Oxford

Human pancreatic islets were isolated from deceased donors under ethical approval obtained from the human research ethics committees in Oxford (REC: 09/H0605/2, NRES committee South Central-Oxford B). All donors gave informed research consent as part of the national organ donation program. Islets were obtained from the Diabetes Research & Wellness Foundation Human Islet Isolation Facility, OCDEM, University of Oxford. All methods and protocols using human pancreatic islets were performed in accordance with the relevant guidelines and regulations in the UK (Human Tissue Authority, HTA). Expression data for *SLC30A8* estimated by RNA sequencing as described previously^57^. For *in vitro* insulin secretion, islets were pre-incubated in Krebs-Ringer buffer (KRB) containing 2 mg/mL BSA and 1 mM glucose for 1 hour at 37°C, followed by 1-hour stimulation in KRB supplemented with 6mM glucose. Insulin content of the supernatant was determined by radioimmunoassay (Millipore UK Ltd, Livingstone, UK) as described previously^58^.

### Human pancreatic islets from LUDC

Human pancreatic islets were obtained from the Human Tissue Laboratory (Lund University, www.exodiab.se/home) in collaboration with The Nordic Network for Clinical Islet Transplantation Program (www.nordicislets.org)^59, 60^. All the islet donors provided their consent for donation of organs for medical research and the procedures were approved by the ethics committee at Lund University (Malmö, Sweden, permit number 2011263). Islet preparation for cadaver donors, their purity check and counting procedure have been described previously^61^. Static i*n vitro* insulin secretion assay from 91 islets (non-diabetic individuals) was performed as described previously^61, 62^. Briefly_,_ six batches of 12 islets per donor were incubated for 1 hour at 37°C in Krebs Ringer bicarbonate (KRB) buffer in presence of 1 mM or 16.7 mM glucose, as well as 1 mM or 16.7 mM glucose together with 70 mM KCl. Insulin concentrations in the extracts was measured using a radioimmunoassay kit (Euro-Diagnostica, Malmö, Sweden). The Association of p.Trp325Arg genotype with expression of *SLC30A8* and other genes involved in insulin production and processing^22^ was performed using RNA sequencing from islets of 140 non-diabetic individuals as described previously^59, 60^. Briefly, RNA sequencing of islets was done using a HiSeq 2000 system (Illumina) for an average depth of 32.4 million paired-end reads (2 × 100 base pairs)^59, 60^. RNA sequencing reads were aligned to hg19 using STAR (Spliced Transcripts Alignment to Reference)^51^. Genome annotations were obtained from the GENCODE (Encyclopedia of Genes and Gene Variants) v20^52^ program and read counting was done using featureCounts^53^. Read counts were normalized to total reads (counts per million) and additionally across-samples normalization was done using TMM method^55^. Association analysis (so called eQTL) was performed on inverse normalized expression values using linear regression adjusted for age, sex and islets purity.

## Statistics

Detail information regarding statistical tests used for each sub-study has been provided in their respective method section or with figure legends.

## Data availability

The data that support the findings of this study are available from the corresponding author on reasonable request. Individual level data for the human study can only be obtained via the Biobank of The Institute of Health and Welfare in Finland.

## URLs

GCTA, http://cnsgenomics.com/software/gcta; SHAPEIT,
http://mathgen.stats.ox.ac.uk/genetics_software/shapeit/shapeit.html, IMPUTE2,
http://mathgen.stats.ox.ac.uk/impute/impute_v2.html;

## Acknowledgements

We thank the Botnia Study Group for recruiting and studying the participants, Jens Juul Holst for measuring GLP-1 concentrations, and Linda Boselli, PhD, for carrying out mathematical modelling of the OGTT studies. We thank Erqian Na for her help with the mouse immunohistochemistry and histology, and Catherine Green and the Chromosome Dynamics & Genome Engineering Cores at the Wellcome Centre for Human Genetics for support with karotyping and genome editing (funded by the Welcome Trust grant 203141). The Botnia and The PPP-Botnia studies (L.G., T.T.) have been financially supported by grants from Folkhälsan Research Foundation, the Sigrid Juselius Foundation, The Academy of Finland (grants no. 263401, 267882, 312063 to LG, 312072 to TT), Nordic Center of Excellence in Disease Genetics, EU (EXGENESIS, EUFP7-MOSAIC FP7-600914), Ollqvist Foundation, Swedish Cultural Foundation in Finland, Finnish Diabetes Research Foundation, Foundation for Life and Health in Finland, Signe and Ane Gyllenberg Foundation, Finnish Medical Society, Paavo Nurmi Foundation, Helsinki University Central Hospital Research Foundation, Perklén Foundation, Närpes Health Care Foundation and Ahokas Foundation, as well as by the Ministry of Education in Finland, Municipal Heath Care Center and Hospital in Jakobstad and Health Care Centers in Vasa, Närpes and Korsholm. The work described in this paper has been supported with funding from collaborative agreements with Pfizer Inc., as well as with Regeneron Genetics Center LLC. J.L. was supported by Vinnova - Sweden’s Innovation Agency (2015-01549), Swedish Diabetes Foundation, Albert Påhlsson Foundation, Hjelt Foundations, Crafoord Foundation, Royal Physiographic Society in Lund, Swedish Foundation for Strategic Research (IRC15-0067), Swedish Research council (2009-1039, Strategic research area Exodiab); E.A. by Crafoord Foundation, Påhlsson Foundation, Swedish Research Council (Dnr: 2017-02688).; O.H. by Lund University Diabetes Center, ALF, Crafoord foundation, Novo Nordisk foundation, Magnus Bergvall foundation, Påhlsson foundation, Diabetes Wellness and Swedish Diabetes Research Foundation; R.C.B. by Italian Ministry of University and Research (PRIN 2015373Z39_004) and University of Parma Research Funds; G.R. by a Wellcome Trust Senior Investigator Award (WT098424AIA), MRC Programme grants (MR/R022259/1, MR/J0003042/1, MR/L020149/1) and Experimental Challenge Grant (DIVA, MR/L02036X/1), MRC (MR/N00275X/1), Diabetes UK (BDA/11/0004210, BDA/15/0005275, BDA 16/0005485) and Imperial Confidence in Concept (ICiC) grants, and a Royal Society Wolfson Research Merit Award. ALG is a Wellcome Trust Senior Fellow in Basic Biomedical Science. M.I.M. and P.R. are Wellcome Senior Investigators. This work was funded in Oxford by the Wellcome Trust (095101 [ALG], 200837 [A.L.G.], 098381 [M.I.M.], 106130 [A.L.G., M.I.M.], 203141 (A.L.G., B.D., M.I.M.], 203141 [M.I.M.], 090531 [P.R.]), Medical Research Council (MR/L020149/1) [M.I.M., A.L.G., P.R.], European Union Horizon 2020 Programme (T2D Systems) [A.L.G.], and NIH (U01-DK105535; U01-DK085545) [M.I.M., A.L.G.]. The research was funded by the National Institute for Health Research (NIHR) Oxford Biomedical Research Centre (BRC) [A.L.G., M.I.M., P.R.]. The views expressed are those of the author(s) and not necessarily those of the NHS, the NIHR or the Department of Health.

## Author Contributions

M.L., L.S., T.T. and L.G. conducted the human study; E.A., O.H., A.B. and J.F. analyzed the genotype data ; M.L., O.P.D., M.T., E.B., R.C.B, T.T. and L.G. analyzed the human data; B.H., N.L.B., S.K.T., M.vD.B., V.C., O.P.D., T.O. and A.L.G. characterized the Human beta-cell model; N.L.B., N.A.J.K., F.A., B.C., D.M., P.K., B.D., M.I.M. and A.L.G. characterized the human IPS cell derived model; U.K., R.P., O.P.D., B.H., A.J.P., I.S., R.R., I.A., P.R., M.I.M. and A.L.G. characterized the human islets; S.K., D.G. and J.G. characterized the *Slc30a8* p.Arg138* mice; D.J., J.L., P.C., A.T., R.C., A-M.R., J.B. and G.R. characterized the rat insulinoma cell-line; M.I.M., A.L.G., T.T. and L.G. supervised the project; O.P.D., M.L., B.H., S.K., N.K., P.R., A.L.G., T.T., and L.G. wrote the manuscript; all authors revised the manuscript.

